# A Zika vaccine based on chimpanzee adenovirus ChAdOx1 elicits lineage-transcending sterile immunity and prevents colonisation of brain and ovaries

**DOI:** 10.1101/514877

**Authors:** César López-Camacho, Stuart Dowall, Victoria Graham, Stephen Findlay-Wilson, Emma Rayner, Young Chan Kim, Roger Hewson, Arturo Reyes-Sandoval

## Introduction

Zika virus (ZIKV) continuous spread has affected more than 80 countries^1^ and no licenced vaccine is currently available^2^. The Asian-lineage is the causative agent of major outbreaks across the globe and most of the current ZIKV vaccine candidates aim to provide homologous efficacy in ZIKV pre-clinical challenge models^2,3^. In contrast, the African-lineage of ZIKV is mainly restricted to sylvatic transmission but not much is known about the ability of this virus to emerge from a sylvatic to human transmission. Efficacy of ZIKV vaccine in mice or macaques relies on the quantification of viraemia by RT-PCR^4^. However, questions remain as to whether current ZIKV vaccine candidates offer lineage-transcending protection and viral clearance in key organs, such as the brain and reproductive tract. Here, we show the cross-protective capabilities of our previously reported simian adenoviral-vectored zika vaccines^5^ (ChAdOx1 ZIKV) against a heterologous African-ZIKV challenge model, in which A129 mice display a high susceptibility to ZIKV infection^6^. Furthermore, we have assessed the ability of our vaccines to prevent infection in tissues such as the brain and ovaries, as both tissues are relevant for ZIKV pathogenesis^7–9^. Finally, we describe a novel ChAdOx1 ZIKV vaccine design carrying a consensus antigen of the African-ZIKV lineage and tested its efficacy in a homologous challenge model.

## Main

The recent Zika outbreaks have emerged from relative obscurity almost 70 years after ZIKV was discovered in Uganda in 1947^10^. Significant outbreaks have been reported in Micronesia^11^ and French Polynesia^12^ in 2007 and 2013-14, respectively; and the largest outbreak was reported in Brazil in early 2015, which spread across the Americas. Phylogenetic studies divide ZIKV into the Asian (^AS^) or African (^AF^)-lineage^13^. The commonality from those outbreaks is that the ZIKV^AS^ was the causal agent. Currently, this lineage still continues to spread to many countries in which the *Aedes* mosquito vector is prevalent, which is an efficient vector of ZIKV. Importantly, ZIKV is a flavivirus known to infect key organs such as the brain^14^ and reproductive tract^15^, causing Gillian Barré syndrome and microcephaly in new-borns. ZIKV has been shown to be sexually-transmitted and to persist long after ZIKV infection in the testis and other fluids such as semen and urine^16^. Therefore, an ideal Zika vaccine must afford complete sterile protection regardless of the ZIKV-lineage and overcome viral persistence in tissues related to the pathogenic features of ZIKV, all in a single and unadjuvanted vaccination approach. While considerable progress has been made at testing several vaccine candidates in Non-Human Primate (NHP) challenge studies^17–20^, in which viraemia is the main correlate of ZIKV infection; little is known of whether the complete elimination of ZIKV from blood can correlate with sterile protection in these immune privileged organs.

Previously, we have reported the generation of four ChAdOx1 ZIKV vaccine candidates based on an Asian-lineage sequence, and their protective efficacy in a homologous ZIKV-lineage challenge model^5^. Although all vaccines demonstrated protective efficacy, we found that a vaccine carrying the prM and the envelope gene without its 3’ transmembrane domain (prME ΔTM) induced long-term anti-envelope antibodies and cleared viraemia with 100% efficacy after only a single and unadjuvanted vaccination^5^. Here, we tested the efficacy of those vaccines against a heterologous ZIKV challenge in a highly susceptible rodent model^6,21^. In addition, we generated a novel ChAdOx1 ZIKV vaccine candidate (ChAdOx1 CprME/NS) based on a consensus sequence from the African-lineage ZIKV (Figure 1a). This carries the full structural gene cassette fused to the non-structural NS2b and NS3 antigens, which are both required to allow the cleavage of Capsid (C) from prM/envelope while exposing the amino terminal region of prM^22^, and tested its efficacy in a homologous ZIKV challenge. 6-8 weeks old, Type-I Interferon receptor knock-out A129 mice were intramuscularly vaccinated (in a blinded experiment) with 10e^8^ IU of ChAdOx1 ZIKV vaccines (n=6), and four weeks after vaccination mice were challenged with a subcutaneous injection of the African-lineage ZIKV strain, MP1751 (Figure 1b). To monitor humoral immune responses elicited by the vaccines, blood samples were taken one day before the ZIKV challenge. After the ZIKV challenge, vaccine efficacy was assessed by measuring the percentage of survival, based on standardised humane time-points such as 20% weight loss and clinical manifestations of ZIKV infection (clinical score)^6^. Animals that met humane clinical endpoints between 7-9 days after challenge (dac) or at the end of the study (21 dac) were culled and blood, spleen, brain, liver and ovaries were collected for histology studies (microscopic lesions), in situ hybridisation (ISH), and to assess ZIKV loads by qRT-PCR.

**Figure 1.**
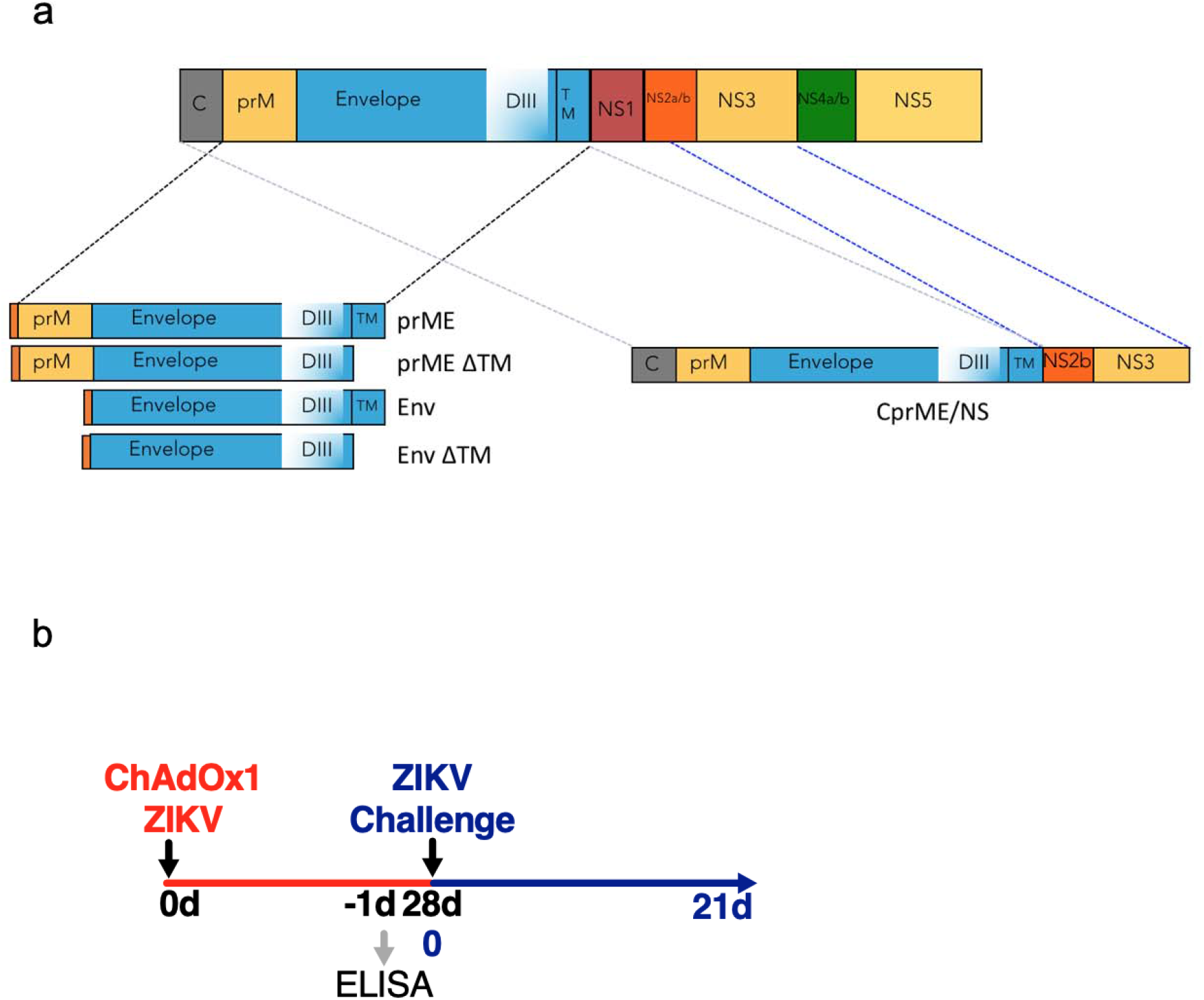
Zika vaccine design. (a) Schematic representation of the Zika immunogen versions used in this study; red box represents the TPA leading sequence. Left diagrams comprise the ZIKV Asian-related vaccines and the right diagram comprise the ZIKV African-related vaccine (b) For ChAdOx1 ZIKV vaccines, A129 mice (n=6 per group) were immunised intramuscularly (i.m.) with a single dose of 10^8^ IU/mice. ZIKV challenge (subcutaneous injection) was performed at day 28 after vaccination using 100 pfu of African strain ZIKV (MP1751). Blood samples were obtained at one day before the ZIKV challenge and at the day animals met humane endpoint or at the end of the study (21 days after challenge).

To gain a comprehensive assessment of vaccine efficacy, temperature-sensing microchips were implanted into A129 mice to monitor the temperature in all vaccinated and control groups. The temperatures were monitored for 30 days before the ZIKV challenge (dbc), including the vaccination day and onwards, and for 21 days after challenge (dac), or until animals reached humane endpoints (Figure 2a and Supplementary Figure 1). Upon challenge, mock-vaccinated (ChAdOx1 Unrelated vaccine) and PBS control mice displayed a gradual increase, followed by a rapid decrease, of temperature (Figure 2a, brown and orange line), which has been described earlier^6^. In contrast, ChAdOx1 ZIKV vaccines were not associated with a gradual increase in temperature (Figure 2a, black arrowhead). The temperatures at the fever stage were significantly lower when compared to control groups (Figure 2b). However, the one-day rapid temperature decrease was observed in the ChAdOx1 Env and ChAdOx1 Env ΔTM groups, which followed the same trend as the unrelated vaccine and PBS groups (Figure 2a, grey arrowhead). ChAdOx1 prME, prME ΔTM and CprME/NS groups did not appear to have this rapid decrease of temperature. After the challenge, all animals were monitored three times a day for clinical signs of disease and a numerical score was assigned for each observation. Clinical signs of disease were observed from 5 to 7 dac in all mice from the unrelated vaccine and the PBS groups (Figure 2c, orange and brown lines). Groups vaccinated with the ChAdOx1 Env presented similar clinical scores as the control groups at d7. A similar trend was observed in a small number of mice from the ChAdOx1 Env ΔTM and milder clinical signs were observed in the ChAdOx1 prME ΔTM and CprME/NS groups (i.e. ruffled fur or hunched). Interestingly, no clinical signs of disease were observed in the ChAdOx1 prME. In parallel, the weight of animals was monitored 30 dbc and 21 dac or until animals met the humane endpoint of 20% of weight loss (Figure 2d). In the control groups, unrelated and PBS, the 20% weight loss occurred first and animals were culled between 6 and 7 dac. ChAdOx1 Env vaccine showed similar weight loss dynamics, although a 2-day delay was observed. The ChAdOx1 Env ΔTM group lost weight rapidly but after 9 dac the animals recovered. ChAdOx1 prME ΔTM and CprME/NS presented <10% weight loss and by the end of the study all the animals recovered normal weight, indicating full recovery. All mice from the ChAdOx1 prME group remained relatively unchanged in the critical days in which control groups lost weight (Figure 2d).

**Figure 2.**
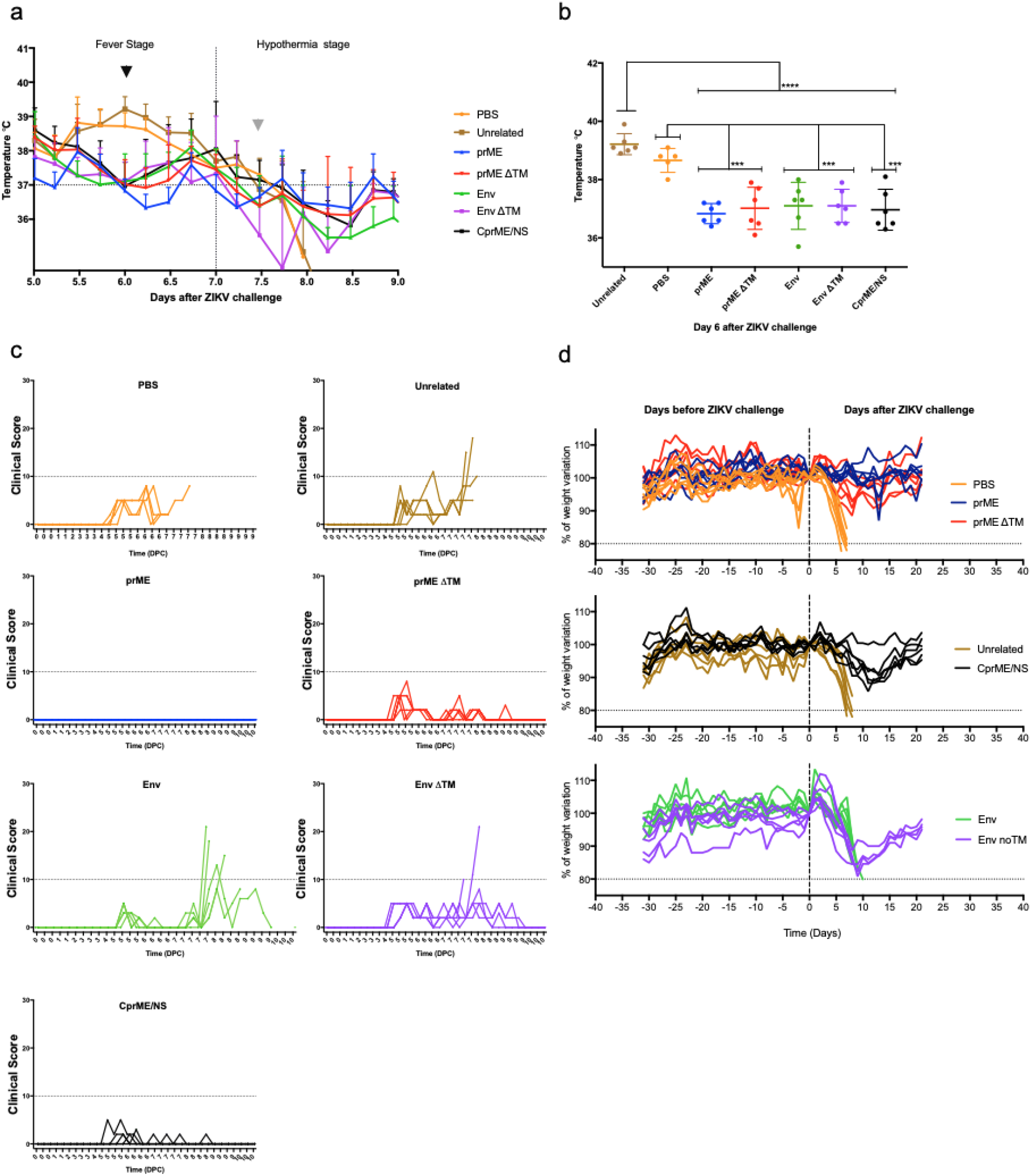
Clinical correlates of protection in A129 mice challenged with ZIKVAF. (a) Difference in temperatures between 5-9 days after ZIKV challenge. Lines represent the mean. (n=6). Complete temperature monitoring from −30 to 21 days after the challenge is shown in Supplementary Figure 1. (b) Mice vaccinated with ChAdOx1 ZIKV vaccines abolished the increase peak of temperature observed at 6 dac in control groups. Lines represent the mean. (n=6). A two-way ANOVA followed by multiple comparison was performed. **** p<0.0001, *** p 0.0003. (c) Clinical manifestation of diseases in control and vaccinated groups. Clinical scores were assigned a numerical value for analysis purposes: 0, normal; 2, ruffled fur; 3, lethargy, pinched, hunched, wasp-waisted; 5, laboured breathing, rapid breathing, inactive, neurological; and 10, immobile. (d) Differences in weight compared to the day of challenge. The weight of all groups of mice were monitored from −30 days to 21 days after challenge.

Based on the humane endpoints (20% weight loss and clinical scores) the percentage of survival was calculated (Figure 3a). Upon challenge with 100 pfu ZIKV, all of the negative control animals met humane clinical endpoints within 7-8 dac. ChAdOx1 Env failed to provide protective efficacy, whilst the deletion of TM from Env in ChAdOx1 Env ΔTM, rescued several animals providing 66% protection. Interestingly, three of the vaccine candidates provided 100% protection: the Asian-lineage based vaccines ChAdOx1 prME and ChAdOx1 prME ΔTM, as well as the African-lineage based vaccine ChAdOx1 CprME/NS. We sought to analyse the levels of anti-ZIKV envelope antibodies at 1 dbc by using ELISA plates coated ZIKV^AS^ envelope^23^ (Figure 3b). Sera from control groups were negative and Log10 reciprocal titers showed that all ChAdOx1 ZIKV vaccines elicited anti-envelope responses. A ZIKV-specific reciprocal of >1 was detected in all ChAdOx1 ZIKV^AS^ vaccinated animals, with the ChAdOx1 prME ΔTM group having the highest titer followed by ChAdOx1 prME. Interestingly, ZIKV envelope antibody titers from the ChAdOx1 CprME/NS sera group were the lowest by ELISA, although all mice from this group were protected after the ZIKV challenge.

**Figure 3.**
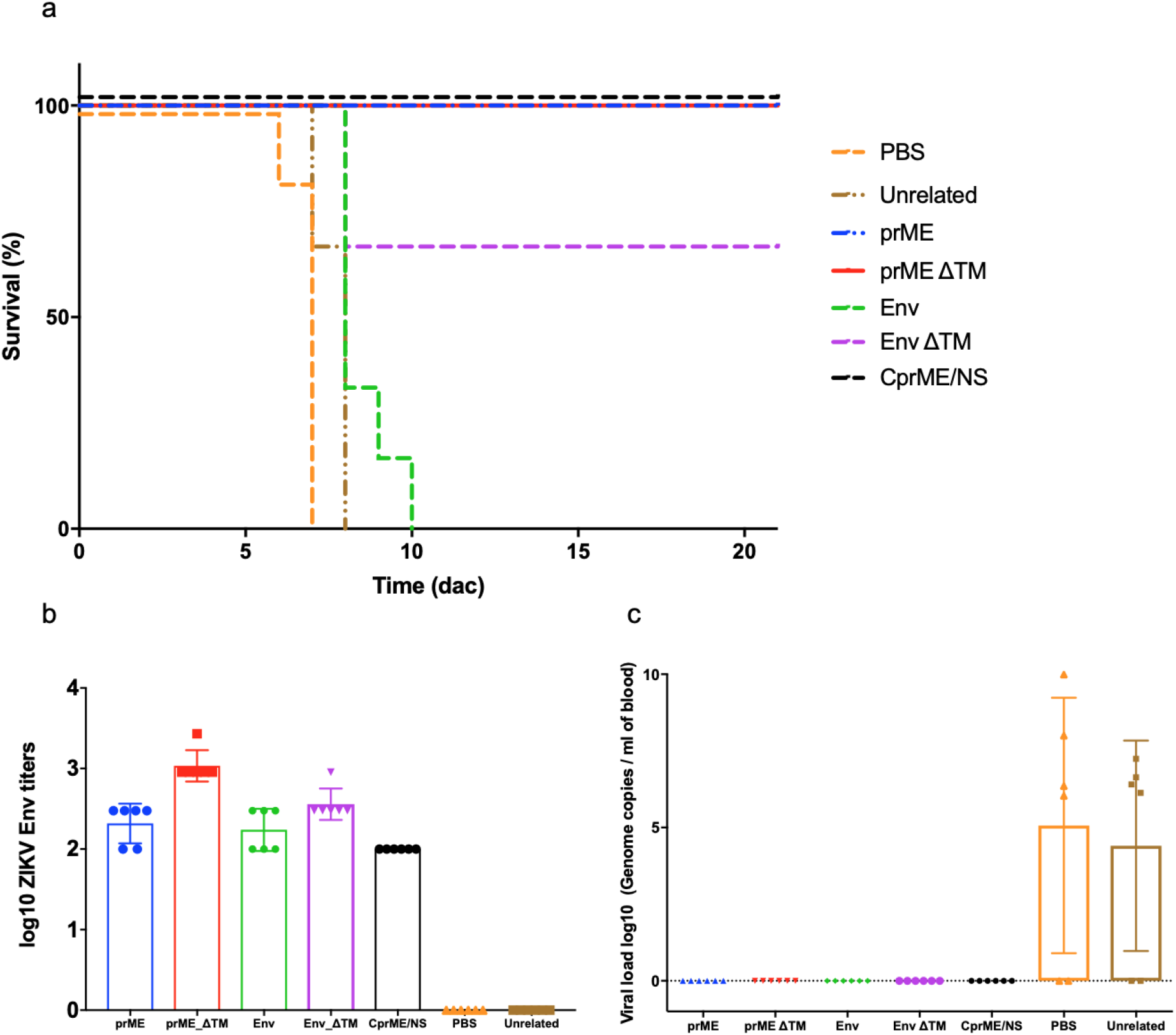
ChAdOx1 vaccine efficacy in protected and unprotected A129 mice. (a) Survival analysis of groups presented as Kaplan-Meier survival curves which shows a range of high, medium and low vaccine efficacy. Lines represent each vaccinated group (n=6). (b) Humoral responses elicited by ChAdOx1 ZIKV vaccines by the time of ZIKV challenge. Antibody responses were quantified by ELISA with plates coated with ZIKV envelope protein. Results are expressed as the log10 of the reciprocal end-point titer. (c) Viral loads in ZIKV challenged groups were monitored in all animals when they met humane endpoints or at the end of the study (21dac) Each dot represents one mouse between each vaccinated or mock control group. Boxes represent the mean and lines represented the standard deviation.

In our previous study^5^ we used a non-lethal challenge model in BALB/C mice and we measured vaccine efficacy *in vivo* by means of partial or complete reduction of viraemia by qRT-PCR. Here, we analysed the viral loads in blood from both protected and unprotected A129 mice by qRT-PCR (Figure 3c). As expected, high levels of viremia were detected in the ChAdOx1 unrelated and the PBS groups. However, each ChadOx1 ZIKV vaccinated group prevented viraemia, regardless of the survival outcome (Figure 3a); i.e. the ChAdOx1 Env group in which all animals met humane endpoints, or the ChAdOx1 prME in which all animals were protected after the ZIKV challenge. These results point out key observations: 1) the presence or absence of viraemia is not a definitive correlate of protective efficacy, and 2) ZIKV, if not found in blood, might be present in different tissues.

To gain a better understanding of protective efficacy versus viral load, we analysed the presence of ZIKV by qRT-PCR in the brain from both the animals that reached humane endpoints and the surviving mice (Figure 4a). For the control groups, unrelated and PBS as well as the ChAdOx1 Env group, ZIKV RNA was detected in the brain at the humane endpoint 7-8 dac. For the vaccinated group ChAdOx1 Env ΔTM (60% survival), viral loads were high in the brain for those animals that were culled by d8, whereas detectable levels where found in 2 out of 4 of surviving animals by 21 dac, which suggests that ZIKV is able to persist long after ZIKV exposure in animals immunised with sub-optimal vaccines. On the other hand, no ZIKV persistence was detected in any of the brain samples from the 100% surviving groups vaccinated with ChAdOx1 prME, and prME ΔTM; and with medium levels of ZIKV in 1 out of 6 animals vaccinated with ChAdOx1 CprME/NS. We assessed the presence of ZIKV by ISH and the histopathological severity associated with ZIKV infection in the brain (Figure 4b, c). As expected, brains from the PBS control group (Figure 4b, PBS micrograph) and from the ChAdOx1 unrelated control group (Supplementary Figure 2a) presented marked inflammation in the meningeal cortex, which correlated with meningeal infiltration by mononuclear inflammatory cells; as well as scattered polymorphonuclear leukocytes (neutrophils) in the parenchyma and perivascular cells plus the presence of scattered, partially degenerated cells in the neurons of the hippocampus (Ammon’s horn), comprising irregularly shaped and partially condensed nuclei (Supplementary Figure 2). Brains from ChAdOx1 Env-vaccinated mice showed similar histopathologic signs as the control groups, such as meningeal infiltration and perivascular cuffing (Figure 4b), which correlated with the presence of viral RNA by ISH (Figure 4c). Similar observations were found in the two animals euthanised at 8 dac from the ChAdOx1 Env ΔTM group, with degeneration of neurons and a large ZIKV presence in the hippocampus (Figure 4c, Env ΔTM) as well as meningeal infiltration, whereas minimal to mild changes in the brain were observed with the remaining animals and staining for the viral antigen was negative. We did not detect marked inflammation of the meninges within the protected groups ChAdOx1 prME, prME ΔTM and CprME/NS. Finally, although the qRT-PCR was negative in the brain samples from groups prME and prME ΔTM, ZIKV RNA was observed in rare cases through single cell-staining by ISH (Figure 4c). However, due to deposits of staining reagent, the presence of rare positive cells might reflect a staining artefact. A summary of the severity of microscopic lesions in the brain is summarised in Supplementary Table 1.

**Figure 4.**
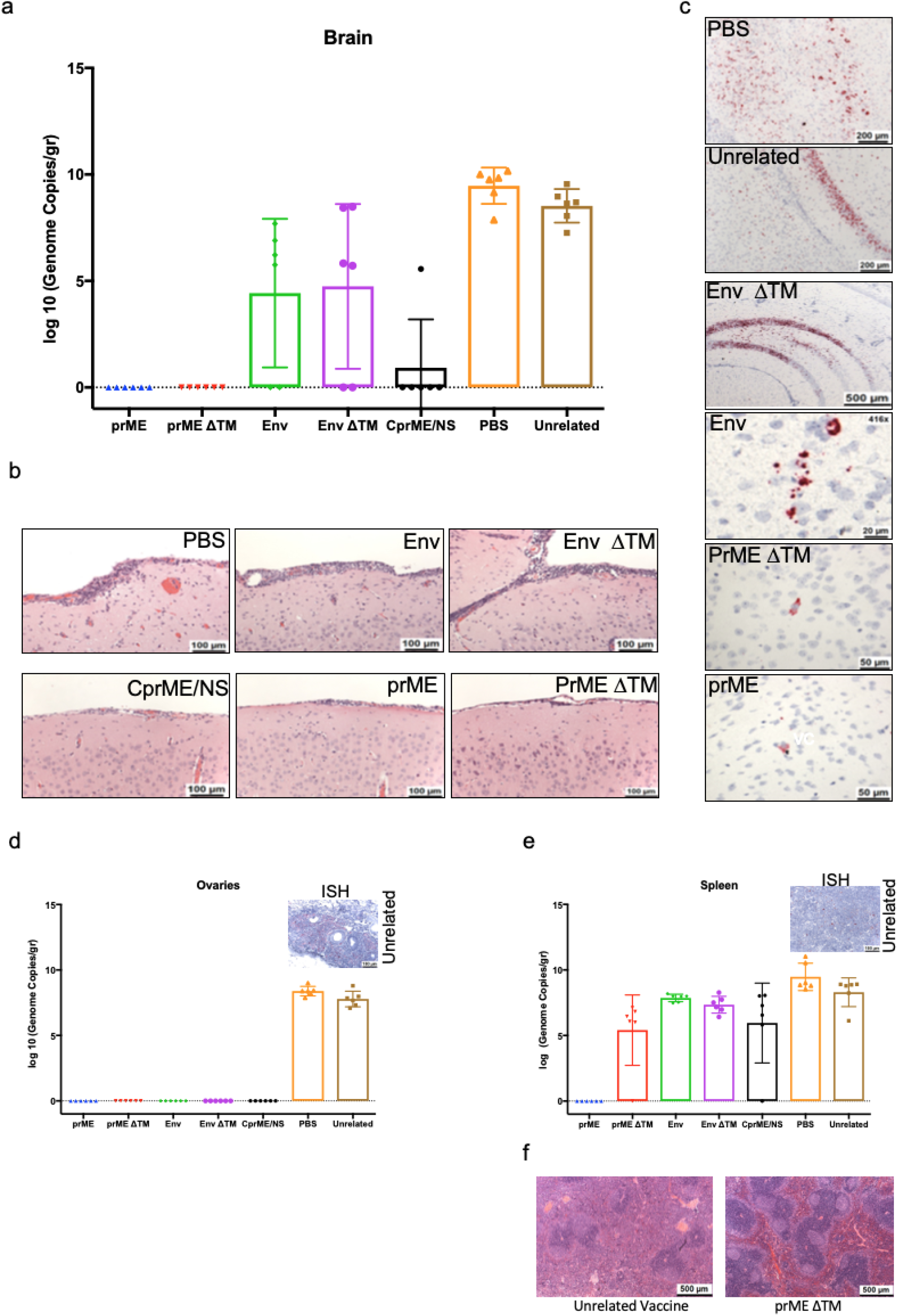
Presence of ZIKV in the brain, ovaries and spleen of A129 mice after ZIKV^AF^ challenge. (a) Levels of ZIKV viral RNA in vaccinated groups at the day of culling or by the end of the study (21 dac), by RT-PCR. Each dot represents one mouse between each vaccinated or mock control group. Boxes represent the mean and lines represented the standard deviation. (b) Pathological findings in the brain of protected and unprotected mice after a ZIKV challenge. Microscopic examination shows inflammation and lymphocyte infiltration of meninges in the control-vaccinated group (PBS) as well as in unprotected mice in both ChAdOx1 Env and ChAdOx1 Env ΔTM groups. The rest of the vaccinated groups did not show meningitis nor lymphocyte infiltration when compared to the control PBS group. Brain tissue was stained with haematoxylin and eosin. Each figure represents a consistent phenotype seen in the rest of each vaccinated group. (c) RNA *in situ* hybridisation analysis to detect ZIKV in the brain of vaccinated and control groups. Red staining reflects ZIKV presence. Strong patchy to diffuse positive staining of viral RNA in the hippocampus (Ammon’s horn) in the control and ChAdOx1 ZIKV vaccine unprotected animals (PBS, ChAdOx1 unrelated vaccine, ChAdOx1 Env and ChAdOx1 Env ΔTM). Very rare staining was detected in mice vaccinated with ChAdOx1 prME and prME ΔTM vaccinated groups. Bar shows scale in microns, accordingly. Levels of ZIKV viral RNA in vaccinated groups at the day of culling or by the end of the study (21 dac), by RT-PCR; in the ovaries (d); and spleen (e). Each dot represents one mouse between each vaccinated or mock control group. Boxes represent the mean and lines represent the standard deviation. ISH for ZIKV RNA presence in ovaries and spleen are presented as a representative example. (f) Demonstrative histology analysis of the spleens from mice vaccinated with the unrelated vaccine and with the ChAdOx1 prME ΔTM.

We extended the analysis to the ovaries in which ZIKV levels were detected in the ChAdOx1 unrelated and PBS groups by qRT-PCR (Figure 4d,). Evidence of infection by ZIKV was reinforced by visualisation of ZIKV RNA by *in situ* hybridisation (ISH) in the unrelated vaccine group, for example (Figure 4d, micrograph). No evidence of ZIKV in the ovary was found in all of the ChAdOx1 ZIKV vaccinated groups assessed by both qRT-PCR and ISH; supporting the clearance of ZIKV in the ovaries of ChAdOx1 ZIKV-vaccinated mice.

Levels of ZIKV in the spleen were also investigated by qRT-PCR. ZIKV RNA was detected in control animals by ISH (Figure 4e). Viral loads were not found in any of the spleens from the ChAdOx1 prME group, whereas 5 out of 6 mice from the ChAdOx1 prME ΔTM group displayed several log fold decreases in viral loads when compared to the groups that received PBS or a ChAdOx1 unrelated vaccine. Importantly, high ZIKV levels were detected in the non-optimal ChAdOx1 Env, the sub-optimal ChAdOx1 Env ΔTM, and in the highly-protective ChAdOx1 CprME/NS.

Histopathological changes in the spleen are primarily considered to be reactive changes to ZIKV infection and provide supportive evidence of infection severity. We found alterations in the spleen in control animals such as poorly defined areas of white pulp (Figure 4f, left panel). This profound alteration was not present in the ChAdOx1 prME, ChAdOx1 prME ΔTM (Figure 4f, right panel) and the ChAdOx1 CprME/NS. Similarly, we found the presence of polymorphonuclear leucocytes (PMN) in the control groups, but normal levels were recorded in all mice vaccinated with the ChAdOx1 prME, prME ΔTM and CprME/NS (Supplementary Table 1). Extra-medullary haematopoiesis (EMH) was observed within all the groups, with marked to moderate effects within the control and ChAdOx1 Env groups, and mild to minimal effects on the rest of ChAdOx1 ZIKV-vaccinated groups (Supplementary Table 1).

Finally, ZIKV RNA was not detected in the liver by either qRT-PCR in any of the groups involved in this study^24^. However, lesions of mild to moderate severity were detected by histology in the control and PBS vaccinated groups, as well as in the ChAdOx1 Env-vaccinated group, while the rest of the ChAdOx1 ZIKV-vaccinated groups presented characteristics within normal limits, with minimal effects in the foci of EMH with apoptosis (Supplementary Table 1).

Taken together, ChAdOx1 ZIKV vaccines demonstrate a range of protective effects upon ZIKV challenge. Adenoviral vectors carrying the full envelope gene of ZIKV but prM showed no protective attributes. Vaccine efficacy is improved leading to partial protection (66%) when the transmembrane domain is deleted (Env ΔTM), and the addition of the prM with envelope increases efficacy to 100 % in the prME, prME ΔTM and CprME/NS vaccine candidates. Of all the vaccine candidates, the ChAdOx1 prME showed the greatest protective correlates, as the animals from this group did not exhibit any signs of disease after ZIKV challenge and viral RNA was not detectable by PCR in any of the tissues collected at necropsy.

The A129 knockout mice lacking the type-I interferon receptor, provide a highly susceptible challenge model for ZIKV^AF^, in which the hallmarks of infectivity are the considerable drop of weight and temperature, together with the clinical signs of infection. Importantly, ZIKV is also able to infect neural and reproductive tissue in this model, and high levels of viremia can be detected in the early days after infection. In this work we have demonstrated the cross-lineage protective efficacy of adenoviral-vectored vaccines expressing the structural antigens of ZIKV^AS^: both ChAdOx1 expressing the prM and Envelope (prME) and the ChAdOx1 prME with a TM deletion (prME ΔTM) afforded 100% survival rate with no massive weight or temperature loss, and no viraemia detected. Here, we report sterile immunity that prevents colonisation of brain and ovaries in female mice, elicited by a single dose of a viral-vectored vaccine. The efficacy of the latter vaccines reflects a very important requirement in ZIKV vaccinology: sterile protection in both neural and reproductive tissue. In humans, ZIKV causes birth defects and neurological conditions such as Gillian Barre syndrome and remains latent within the sexual organs.

In this study, we vaccinated and challenged female A129 mice with ZIKV, it would be very useful to follow up the efficacy of this and other vaccine candidates in gender-randomised groups of mice, to assess the presence of ZIKV in reproductive tissue such as testis, ovaries, and the reproductive tract. More importantly, testing those vaccines in a ZIKV-transmission model would be essential to completely rule-out ZIKV transmission through the sexual route.

In this work, we found that non-protected vaccinated mice (i.e. ChAdOx1 Env) met humane endpoints by eight dac, and surprisingly, no ZIKV was detected in the blood of these animals by RT-PCR, but high levels were detected in the brain. Furthermore, 66% of the mice vaccinated with ChAdOx1 Env ΔTM showed full recovery after challenge, and yet we were still able to detect viral within the brain 21 dac, suggesting that ZIKV persistence is an important feature worthy of further clinical and preclinical vaccine developments. Equally important is the fact that ZIKV challenge models in NHP rely mainly on the presence or absence of ZIKV in the blood. Interestingly, ZIKV persistence in NHP has been documented in the cerebrospinal fluid and lymph nodes^25^. It is important to acknowledge that although viraemia is an important correlate of vaccine efficacy, this parameter may not reflect the capacity of other vaccine candidates to generate protection in other organs such as neural and reproductive tissue.

Finally, we have developed a Zika vaccine based on the African ZIKV-lineage, which contains the whole structural cassette and fused to the non-structural proteins NS2b/NS3. This vaccine conferred 100% protective efficacy as none of the animals met humane end-points. However, the average of ELISA endpoint titers to ZIKV envelope was low in this group in comparison to those from mice protected or unprotected (ChAdOx1 prME ΔTM and ChAdOx1 Env, respectively). Furthermore, ZIKV RNA was detected in the brain of 1 out of 6 animals at 21 dac and we cannot rule out whether this is viral RNA traces nor it has infective capabilities. The question remains whether these observations are due to a suboptimal elicitation of immune responses, or a suboptimal vaccination with ChAdOx1 CprME/NS. Finally, ChAdOx1CprME/NS contains the antigen NS2b and importantly NS3, which has been shown to induce T cell immunity in ZIKV, DENV and other flavivirus^26^. Future studies comparing CprME/NS versus the sole insertion of the ZIKV NS3 will provide further insights into the mode of action of this vaccine. Harnessing ZIKV vaccines that are able to elicit both antibodies against structural antigens and specific T cell responses against non-structural proteins or vice versa, create new and interesting possibilities in the search for suitable ZIKV vaccine candidates.

## Material and Methods

### Animals

Forty-two female A129 mice (5-6 weeks old) were purchased from a home office approved breeder and supplier (B…K Universal Ltd, part of Marshall BioResources). The mice were implanted with a temperature and identity chip (idENTICHIP with Bio Thermal) upon arrival (n=6 mice per group). During three-day rest, baseline observations of behaviour, temperature and weight were recorded. The experimental design accounted for the 3R reduction (Replacement, Reduction, Refinement). No randomisation was used in this work.

### Transgene design

Design and production of the ZIKV Asian-based vaccines ChAdOx1 Env, ChAdOx1 Env ΔTM, ChAdOx1 prME and ChAdOx1 prME ΔTM have been previously reported^5^. For the African Lineage vaccine design, DNA coding the Capsid (C), prM and Envelope (E) coding sequences (794 a.a.), joined to the NS2b (127 a.a.) and NS3 (616 a.a.) was codon optimised. Transgene design included the required enzymatic restriction sites to allow the in-frame cloning of the transgene between the CMV promoter and the Poly(A) sequence region contained in our shuttle and expression vector (pMono). Synthetic gene cassette was produced by GeneArt and was named CprME/NS.

### Vaccine production of ChAdOx1 CprME/NS

The CprME/NS transgene was digested with KpnI and NotI to allow in-frame ligation between the CMV promoter and the Poly(A) regions contained into an expression plasmid (pMono). pMono CprME/NS plasmid was expanded in E. coli. The entry plasmid containing attL regions sequences was recombined with those attR regions contained in the destination vector ChAdOx1 using an *in vitro* Gateway reaction (LR Clonase II system, Invitrogen). Successfully recombined ChAdOx1 CprME/NS plasmid was verified by DNA sequencing using flanking primers (promoter forward primer and Poly-(A) Reverse primer). Standard cell biology and virology techniques were performed to generate the non-replicative adenoviral vectors^27^.

### Immunisation of mice

Mice were vaccinated with a single dose of chimpanzee adenoviral vectored vaccine (ChAdOx1) encoding the ZIKV antigens at a dose of 1×10^8^ infectious units (IU). All vaccines were administered intramuscularly (i.m.) in the right hind limb and diluted in endotoxin-free PBS. ChAdOx1 expressing an unrelated malarial antigen (cTRAP) were used as control vaccines.

### Ethics statement

All animals and procedures were used in accordance with the terms of the UK Home Office Animals Act Project License. Animal experiments performed at PHE were approved

These studies were approved by the ethical review process of Public Health England, Porton Down, UK, and by the Home Office, UK via an Establishment Licence (PEL PCD 70/1707) and project licence (30/3147). A set of humane endpoints based on clinical manifestation of disease were defined in the protocol of the project licence and are described below.

### Challenge virus

All mice were challenged with 100 pfu of African strain ZIKV (MP1751) via the subcutaneous (s.c.) route to mimic mosquito bite; 40ul into the right leg and 40l into left leg toward the ankle.

### Sample collection

When animals met humane clinical endpoints or at the end of the study (21 days post-challenge) they were culled by an approved Schedule 1 method. The following samples were collected at necropsy:

Blood: 100 μl in an RNA protect tube and frozen in an ultra-low freezer with the reminder into serum separation tubes.

Spleen: half to be placed in Precellys tubes and frozen in an ultra-low freezer, the remainder in a histology pot

Brain: 2/3rds LHS+ (or most intact) to be placed in a histology pot, the remainder to be placed in Precellys tubes and frozen in an ultra-low freezer.

Liver: small section to be placed in Precellys tubes and frozen in an ultra-low freezer. A section (no more than 3mm thick) to be placed into a histology pot

Ovaries: LHS to be placed in a histology pot, the RHS to be placed in Precellys tubes and frozen in an ultra-low freezer.

### Clinical Measurements

During the course of the study, weights and temperatures were collected at least once daily and clinical scores assessed at least twice a day. During critical periods of the study the monitoring intensity was increased to allow humane clinical endpoints to be more rapidly captured and reduce any potential suffering of animals with severe disease. Clinical scores were assigned a numerical value for analysis purposes: 0, normal; 2, ruffled fur; 3, lethargy, pinched, hunched, wasp-waisted; 5, laboured breathing, rapid breathing, inactive, neurological; and 10, immobile. To prevent unnecessary suffering to animals, humane clinical endpoints were used where animals would be culled upon reaching any of the following criteria:

Immobility defined as a lack of movement even after a stimulus such as handling; Neurological signs including repetitive or unusual movement; Weight loss of 20% or more compared to maximum recorded weight from days 1-3 pre-challenge or prevaccination; or continual vocalisation.

### Pre-challenge bleed

On day 27 post-vaccination, a blood sample from each mouse was taken (no more than 100 μl) into SST tubes for serum isolation.

### Whole IgG Enzyme-linked immunosorbent assay

96-well plates coated with ZIKV envelope protein and blocked with Pierce Blocking Protein-free reagent. Serial dilutions (3-fold) of sera from vaccinated mice were added. Diluted sera were incubated at RT for 1 Lh, and after four times washing buffer incubations, 100 ☐μl/well of anti-mouse IgG HRP-conjugate working solution was added for 2 hr at room temperature. Plates were washed 5 times and developed 15☐min at room temperature with 100☐μl of the developing substrate, then stopped by the addition of 100☐μl of stop solution (NaOH 1M). Absorbance was measured at 405☐nm on a microplate reader. ELISA ODs were compared between all vaccinated groups at different sera dilutions. For pre-challenge ELISA, endpoint titers were defined as the highest reciprocal serum dilution that yielded an absorbance >2-fold over background values.

### Assessment of viral load by PCR

Tissue samples were weighed and homogenised with PBS using ceramic beads and an automated homogeniser (Precellys, UK) with a programme consisting of six 5 second cycles of 4500 rpm with a 30 second gap. 100 μl of tissue homogenate was transferred to 300 μl RLT buffer (Qiagen, UK) for RNA extraction; for blood 25 μl from the RNAprotect tubes and 75 μl of PBS was to 300 μl of RLT buffer. After at least 10 minutes, 400 μl of 70% ethanol was added to the blood or tissue homogenate / RLT. The sample underwent further processing by passing through a Qiashredder (Qiagen, UK). The RNA was isolated from liver, ovary and brain using the Biosprint extraction kit (Qiagen, UK) and the Kingfisher flex system (ThermoFisher Scientific, UK). For the spleen and blood samples, RNA was manually extracted using an RNeasy mini kit (Qiagen, UK).

A ZIKV specific real-time RT-PCR assay was utilised for the detection of viral RNA with primer and probe sequences adopted from a published method with in-house optimisation and validation performed to provide optimal mastermix and cycling conditions. Real-time RT-PCR was performed using the SuperScript III Platinum One-step qRT-PCR kit (Life Technologies, UK). The final mastermix (15 μl) comprised 10 μl of 2x Reaction Mix, 1.2 μl of PCR-grade water, 0.2 μl of 50 mM MgSO4, 1 μl of each primer (ZIKV 1086 and ZIKV 1162c both at 18 μM working concentration), 0.8 μl of probe (ZIKV 1107-FAM at 25 μM working concentration) and 0.8 μl of SSIII enzyme mix. 5 μl of template RNA was added to the mastermix in order to give a final reaction volume of 20 μl. The cycling conditions used were 50°C for 10 minutes, 95°C for 2 minutes, followed by 45 cycles of 95°C for 10 seconds and 60°C for 40 seconds with a final cooling step of 40°C for 30 seconds. Quantification analysis using fluorescence was performed at the end of each 60°C step. Reactions were run and analysed on the QuantStudio platform (Applied Biosystems, UK).

Quantification of viral load in samples was performed using a dilution series of quantified RNA oligonucleotide (Integrated DNA Technologies). The oligonucleotide comprised the 77 bases of ZIKV RNA targeted by the assay, based on GenBank accession AY632535.2 and was synthesised to a scale of 250 nmol with HPLC purification.

### Histology

Samples of brain, ovary, liver and spleen were fixed in 10% neutral buffered saline and processed routinely to paraffin wax. Sections were cut at 3-5 μm, stained with haematoxylin and eosin and examined microscopically. Lesions referable to infection were scored subjectively using the following scale: within normal limits, minimal, moderate and marked. The pathologist was blinded to the groups in order to prevent bias.

### RNA in situ hybridisation (ISH)

RNA ISH was performed with an RNAscope 2.5 (Advanced Cell Diagnostics) according to the manufacturer’s instructions. In brief, formalin-fixed paraffin-embedded tissue sections were deparaffinised by incubation at 60 min at 60°C. Hydrogen peroxide treatment for 10 min at room temperature quenched endogenous peroxidases. Slides were then boiled for 15 min in RNAscope Target Retrieval Reagents and incubated for 30 min in RNAscope Protease Plus before hybridisation. For probes, V-ZIKA-pp-O1-sense (Advanced Cell Diagnostics, catalogue no. 463791) was used due to having 99% specificity with the MP1751 challenge strain of ZIKV. Tissues were counterstained with Gill’s haematoxylin and visualised with standard bright-field microscopy. Each slide was scanned systematically so all areas of the tissue were assessed.

### Statistics

Graphing and statistical analysis were performed using GraphPad Prism version 7.00 for Mac, GraphPad Software, La Jolla California USA, www.graphpad.com. To assess differences between groups, a two-way ANOVA followed by Multiple comparison was performed. All graphs were plotted with bars and errors, representing the mean and the s.d. Dots represent each individual between groups. Assumptions of data are: normal distribution, homogeneity, independent, and groups must have same sample size.

### Data availability

The authors declare that the data supporting the findings of this study are shown in the article and in the Supplementary section and available from the authors upon request.

## Acknowledgements

We would like to thank the Jenner Institute’s Vector Core Facility for producing and supplying the recombinant viral vectors and the Biological Investigations Group at Public Health England for the care of animals. A. R-S is a Jenner Investigator and an Oxford Martin Fellow. This report is work commissioned by Innovate UK and the UK Department of Health and Social Care (projects No. 972216 and 971557). The views expressed in this publication are those of the author(s) and not necessarily those of Innovate UK or the Department of Health and Social Care.

## Contributions

A.R.S. is the grant holder and directed the project and commissioned the work. C.L.C. designed, constructed, characterised the vaccines and analysed the data. S.D., V.G. and R.H. conceived and designed the animal experiments. S.D., V.G., E.R. and S.F.W. performed the animal experiments including the ZIKV challenge model and the RT-PCR viral loads, microscopy and histology. C.L.C. performed ELISA assays and analysed ZIKV challenge data. Y.C.K. assisted with ELISA experiments and produced the ZIKV envelope protein. All authors read and commented the manuscript. C.L.C. and A.R.S. wrote the manuscript.

## Competing interests

A.R.S. and C.L.C. are co-inventors of the Zika vaccines described in this manuscript, filed by Oxford University Innovation Limited in the International Patent Application No. PCT/GB2017/052220 Zika Vaccine, A.V.S.H. and S.C.G. are co-inventors on a patent application (WO/2012/172277) on the ChAdOx1 viral vector filed by Oxford University Innovation. All other co-authors declare neither financial nor non-financial competing interests.

---

**Supplementary Figure 1.**
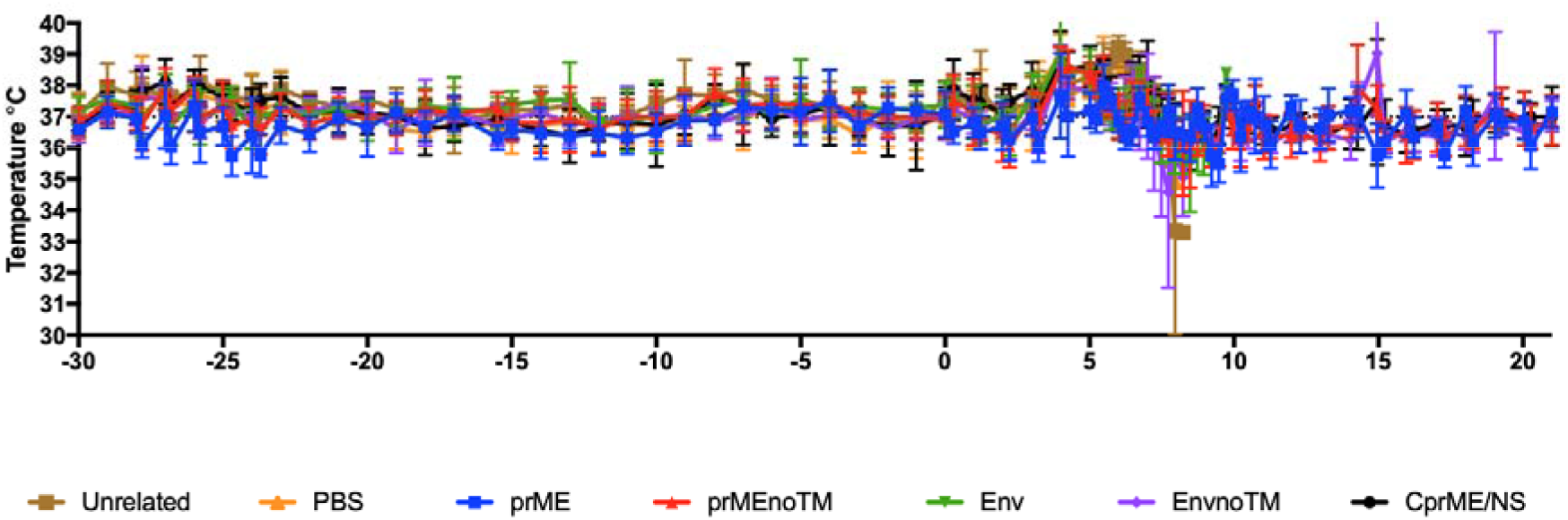
Temperature monitoring from 30 dbc and 21 dac. A temperature sensing chip was implanted in each mouse involved in the study. Temperature monitoring was mainly done twice a day. Lines represent the mean of each vaccinated group (n=6). Bar errors represent the standard deviation for each time point.

**Supplementary Figure 2.**
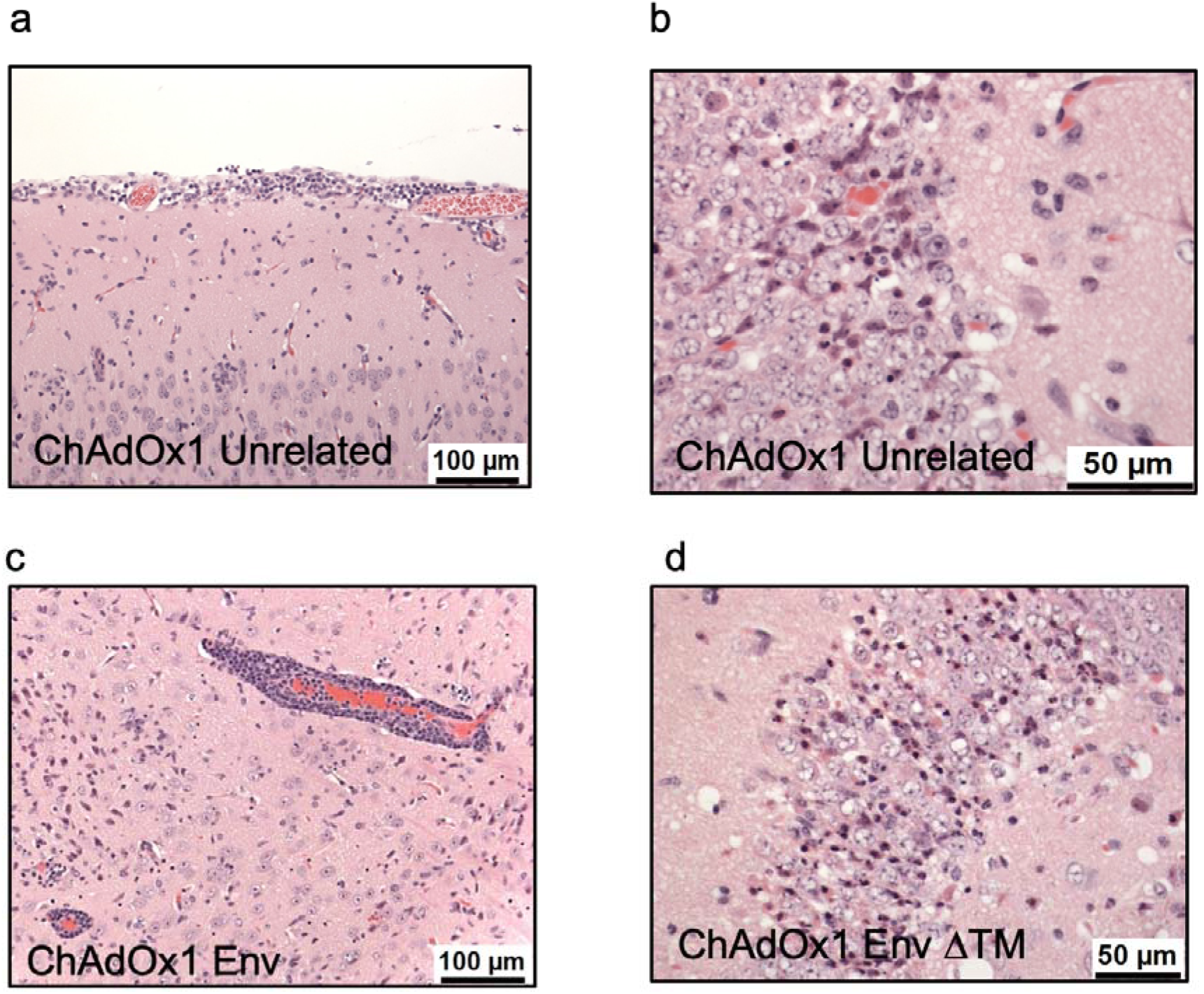
Pathological findings monitored by brain histology in vaccinated-A129 mice after ZIKV challenge. (a) Brain, meninges. Animal 13690, 727/17, (ChAdOx1 pvTRAP-Unrelated). Infiltration of mainly mononuclear cells, mainly mononuclear into the meninges HE. (b) Brain, hippocampus. Animal 34144, 728/17, (ChAdOx1 pvTRAP- Unrelated). Diffuse cellular degeneration in Ammon’s horn. Degenerating neurons are observed (intense dark staining). Presence of scattered, partially degenerated cells in the neurons of the hippocampus (Ammon’s horn), comprising hyper-eosinophilic cytoplasm and irregularly, shaped, partially condensed nuclei. (c) Brain. Animal 31522, 707/16, (ChAdOx1 Env). Variable, perivascular cuffing of blood vessels in the parenchyma and meninges by mononuclear cells, many having the morphology of lymphocytes. (d) Brain. Animal 12472, 708/17. Moderate degeneration of neurons in the hippocampus. HE. In this group, changes associated with ZKV infection in the brain were noted variably in all animals ranging from minimal to moderate. The most severe changes were noted in unprotected animals 12472 and 13688, both of which were euthanised early for welfare reasons at 8 dac.

**Supplementary Table 1.**
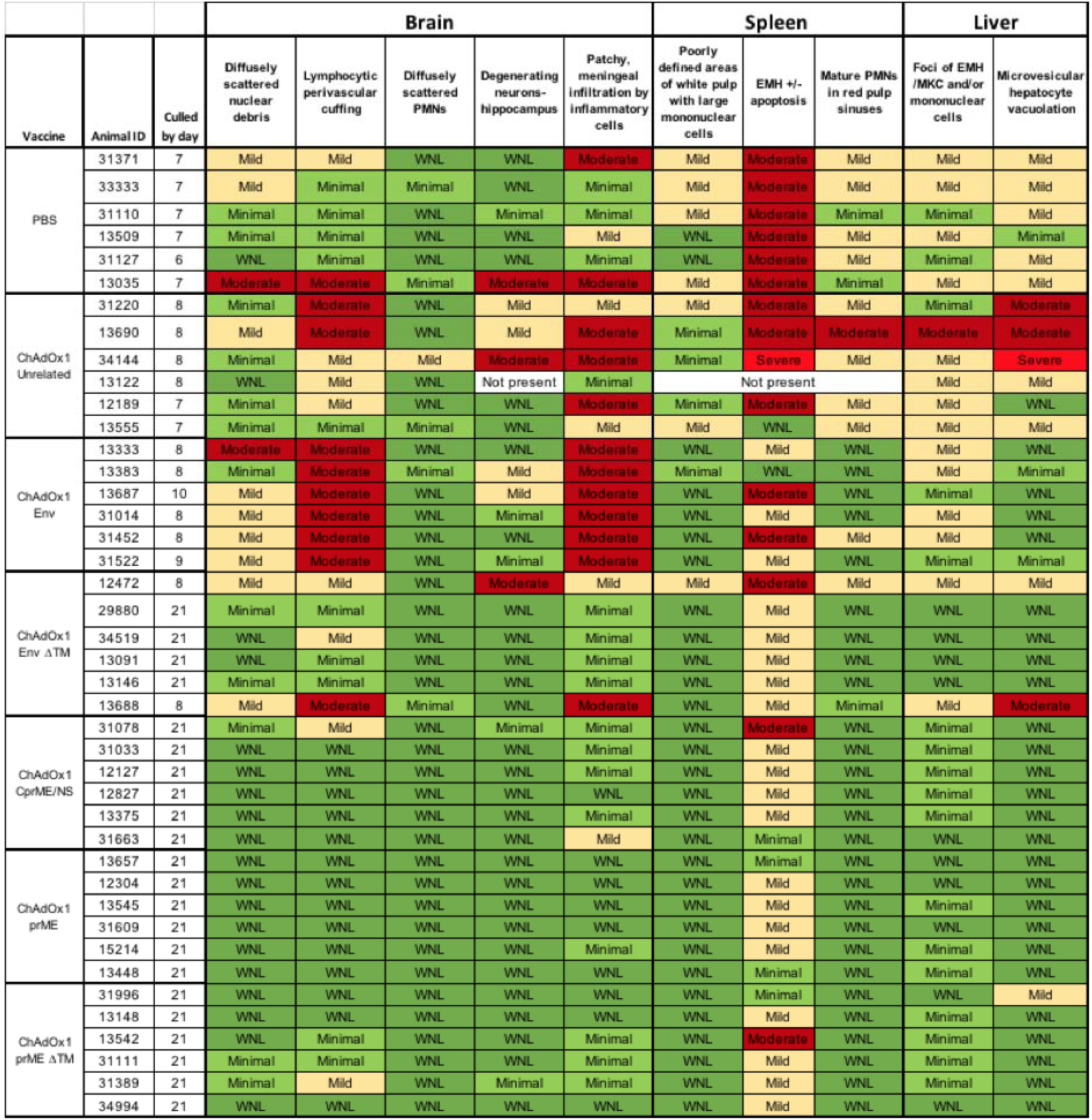
Histology changes observed in A129 mice challenged with ZIKV. Table shows changes in Brain, Spleen and Liver. EMH, extramedullary apoptosis. PMNs, polymorphonuclear lymphocytes. WNL, within normal limits. Colours depict severity of lesions, being green minimal or WNM, yellow being mild and red being moderate to severe.

| Vaccine | Animal ID | Culled by day | Brain |  |  |  |  | Spleen |  |  | Liver |  |
| --- | --- | --- | --- | --- | --- | --- | --- | --- | --- | --- | --- | --- |
|  |  |  | Diffusely scattered nuclear debris | Lymphocytic perivascular cuffing | Diffusely scattered PMNs | Degenerating neurons-hippocampus | Patchy, meningeal infiltration by inflammatory cells | Poorly defined areas of white pulp with large mononuclear cells | EMH +/- apoptosis | Mature PMNs in red pulp sinuses | Foci of EMH /MKC and/or mononuclear cells | Microvesicular hepatocyte vacuolation |
| PBS | 31371 | 7 | Mild | Mild | WNL | WNL | Moderate | Mild | Moderate | Mild | Mild | Mild |
|  | 33333 | 7 | Mild | Minimal | Minimal | WNL | Minimal | Mild | Moderate | Mild | Mild | Mild |
|  | 31110 | 7 | Minimal | Minimal | WNL | Minimal | Minimal | Mild | Moderate | Minimal | Minimal | Mild |
|  | 13509 | 7 | Minimal | Minimal | WNL | WNL | Mild | WNL | Moderate | Mild | Mild | Minimal |
|  | 31127 | 6 | WNL | Minimal | WNL | WNL | Minimal | WNL | Moderate | Mild | Minimal | Mild |
|  | 13035 | 7 | Moderate | Moderate | Minimal | Moderate | Moderate | Mild | Moderate | Minimal | Mild | Mild |
| ChAdOx1 Unrelated | 31220 | 8 | Minimal | Moderate | WNL | Mild | Mild | Mild | Moderate | Mild | Minimal | Moderate |
|  | 13690 | 8 | Mild | Moderate | WNL | Mild | Moderate | Minimal | Moderate | Moderate | Moderate | Moderate |
|  | 34144 | 8 | Minimal | Mild | Mild | Moderate | Moderate | Minimal | Severe | Mild | Mild | Severe |
|  | 13122 | 8 | WNL | Mild | WNL | Not present | Minimal | Not present |  |  | Mild | Mild |
|  | 12189 | 7 | Minimal | Mild | WNL | WNL | Moderate | Minimal | Moderate | Mild | Mild | WNL |
|  | 13555 | 7 | Minimal | Minimal | Minimal | WNL | Mild | Mild | WNL | Mild | Mild | Mild |
| ChAdOx1 Env | 13333 | 8 | Moderate | Moderate | WNL | WNL | Moderate | WNL | Mild | WNL | Mild | WNL |
|  | 13383 | 8 | Minimal | Moderate | Minimal | Mild | Moderate | Minimal | WNL | WNL | Mild | Minimal |
|  | 13687 | 10 | Mild | Moderate | WNL | Mild | Moderate | WNL | Moderate | WNL | Minimal | WNL |
|  | 31014 | 8 | Mild | Moderate | WNL | Minimal | Moderate | WNL | Mild | WNL | Mild | WNL |
|  | 31452 | 8 | Mild | Moderate | WNL | WNL | Moderate | WNL | Moderate | Mild | Mild | WNL |
|  | 31522 | 9 | Mild | Moderate | WNL | Minimal | Moderate | WNL | Mild | WNL | Minimal | Minimal |
| ChAdOx1 Env ΔTM | 12472 | 8 | Mild | Mild | WNL | Moderate | Mild | Mild | Moderate | Mild | Mild | Mild |
|  | 29880 | 21 | Minimal | Minimal | WNL | WNL | Minimal | WNL | Mild | WNL | WNL | WNL |
|  | 34519 | 21 | WNL | Mild | WNL | WNL | Minimal | WNL | Mild | WNL | WNL | WNL |
|  | 13091 | 21 | WNL | Minimal | WNL | WNL | Minimal | WNL | Mild | WNL | WNL | WNL |
|  | 13146 | 21 | Minimal | Minimal | WNL | WNL | Minimal | WNL | Mild | WNL | WNL | WNL |
|  | 13688 | 8 | Mild | Moderate | Minimal | WNL | Moderate | WNL | Mild | Minimal | Mild | Moderate |
| ChAdOx1 CprME/NS | 31078 | 21 | Minimal | Mild | WNL | Minimal | Minimal | WNL | Moderate | WNL | Minimal | WNL |
|  | 31033 | 21 | WNL | WNL | WNL | WNL | Minimal | WNL | Mild | WNL | Minimal | WNL |
|  | 12127 | 21 | WNL | WNL | WNL | WNL | Minimal | WNL | Mild | WNL | Minimal | WNL |
|  | 12827 | 21 | WNL | WNL | WNL | WNL | WNL | WNL | Mild | WNL | Minimal | WNL |
|  | 13375 | 21 | WNL | WNL | WNL | WNL | Minimal | WNL | Mild | WNL | Minimal | WNL |
|  | 31663 | 21 | WNL | WNL | WNL | WNL | Mild | WNL | Minimal | WNL | WNL | WNL |
| ChAdOx1 prME | 13657 | 21 | WNL | WNL | WNL | WNL | WNL | WNL | Minimal | WNL | WNL | WNL |
|  | 12304 | 21 | WNL | WNL | WNL | WNL | WNL | WNL | Mild | WNL | WNL | WNL |
|  | 13545 | 21 | WNL | WNL | WNL | WNL | WNL | WNL | Mild | WNL | Minimal | WNL |
|  | 31609 | 21 | WNL | WNL | WNL | WNL | WNL | WNL | Mild | WNL | WNL | WNL |
|  | 15214 | 21 | WNL | WNL | WNL | WNL | Minimal | WNL | Mild | WNL | Minimal | WNL |
|  | 13448 | 21 | WNL | WNL | WNL | WNL | WNL | WNL | Minimal | WNL | Minimal | WNL |
| ChAdOx1 prME ΔTM | 31996 | 21 | WNL | WNL | WNL | WNL | WNL | WNL | Minimal | WNL | WNL | Mild |
|  | 13148 | 21 | WNL | WNL | WNL | WNL | WNL | WNL | Mild | WNL | Minimal | WNL |
|  | 13542 | 21 | WNL | Minimal | WNL | WNL | Minimal | WNL | Moderate | WNL | Minimal | WNL |
|  | 31111 | 21 | Minimal | Minimal | WNL | WNL | Minimal | WNL | Mild | WNL | Minimal | WNL |
|  | 31389 | 21 | Minimal | Mild | WNL | Minimal | Minimal | WNL | Mild | WNL | Minimal | WNL |
|  | 34994 | 21 | WNL | WNL | WNL | WNL | WNL | WNL | Mild | WNL | WNL | WNL |

